# Long non-coding RNA L13Rik promotes high glucose-induced mesangial cell injury by regulating miR-2861/CDKN1B axis

**DOI:** 10.1101/2022.06.02.494486

**Authors:** Linlin Sun, Miao Ding, Fuhua Chen, Dingyu Zhu, Xinmiao Xie

**Affiliations:** Department of Nephrology, Tongren Hospital, Shanghai Jiao Tong University School of Medicine, Shanghai, China

**Keywords:** Diabetic nephropathy, mesangial cell, L13Rik, miR-2861, CDKN1B

## Abstract

Diabetic nephropathy (DN) is a frequent and severe microvascular complication of diabetes. Glomerular mesangial cell (MC) injury occurs at the initial phase of DN and acts as a critical role in the pathogenesis of DN. Given the importance of long non-coding RNA (lncRNA) in regulating MC hyperplasia and extracellular matrix (ECM) accumulation, it is essential to identify functional lncRNAs during MC injury. Here a novel lncRNA, C920021L13Rik (L13Rik for short), was identified to up-regulated in DN progression. The expression of L13Rik in DN patients and diabetic rats was assessed using quantitative real-time PCR (qRT-PCR), and the function of L13Rik on regulating HG-induced MC injury was assessed using cell counting kit-8 (CCK-8) and western blot assay to analyze MC viability and ECM accumulation. We found that L13Rik level was significantly increased while miR-2861 level was significantly decreased in peripheral blood of DN patients, renal tissues of diabetic rats, and HG-treated MCs. Functionally, both L13Rik depletion and miR-2861 overexpression effectively reduced HG-induced MC survival, ECM accumulation, and cell hypertrophy. Mechanistically, L13Rik functioned as a competing endogenous RNA (ceRNA) to sponge miR-2861, resulting in the de-repression of its target cyclin-dependent kinase inhibitor 1B (CDKN1B), a gene known to accelerate MC injury. Collectively, the current results demonstrate that up-regulated L13Rik is correlated with DN, and may be a hopeful therapeutic target for DN.

## 1. Introduction

Diabetic nephropathy (DN), happens in as much as 50% of living diabetes, is a severe diabetic microvascular complication and a major cause of chronic kidney disease (CKD) and end-stage renal disease worldwide (Alicic et al., 2017, Selby and Taal, 2020). At the early stage of DN, MCs proliferate abnormally and result in the excessive ECM accumulation, which are two critical pathological characteristics of CKD (Rojas-Canales et al., 2019, Sun, 2019). Currently, managing blood sugar and blood pressure and blockading renin-angiotensin-aldosterone system (RAAS) are conventional treatment for retarding DN progression (Choudhury et al., 2010, Chang et al., 2020). However, there is no effective treatment to stop or reverse that. It is essential to reveal the mechanisms by which HG triggers MC proliferation and excessive ECM accumulation.

With recent advances in RNA sequencing and bioinformatics algorithms, long non-coding RNAs (lncRNAs) are no longer considered as “transcriptional noise” because they do not have protein-coding potential (Okazaki et al., 2002, Jiang and Zhang, 2021). More and more studies have demonstrated that lncRNA exerts a crucial role in various kinds of physiopathological processes such as genomic imprinting, gene expression, cell proliferation, tumorigenesis (de Oliveira et al., 2019, Bach and Lee, 2018, Sakatani et al., 2005). As expected, lncRNAs are abnormally expressed in DN patients and mice and exert an important role in the progression of DN (Coellar et al., 2021). The data from RNA sequencing show that 106 lncRNAs are dys-regulated in kidney tissues of DN mice and 42 lncRNAs are correlated with DN (Li et al., 2018). Functional experiments demonstrated that down-regulated lncRNA-1700020I14Rik attenuates MC proliferation and subsequent ECM accumulation (Li et al., 2018). Zhang et al. showed that 95 lncRNAs are dys-expressed in kidney tissues of DN mice compared with normal mice and up-regulated lncRNA-Rpph1 accelerates MC proliferation and inflammatory response through interacting with DN-related factor galectin-3 (Zhang et al., 2019). LncRNA-CDKN2B-AS1 expression is increased in peripheral blood of DN patients and HG-treated MCs, and CDKN2B-AS1 depletion contributes to alleviate MC proliferation and ECM accumulation by interacting with miR-424-5p (Li et al., 2020). Given the importance of lncRNA in DN, it is essential to identify functional lncRNAs during MC injury. L13Rik is an up-regulated lncRNA in renal tissues from type 2 diabetes mice compared with normal mice (GSE642 dataset, https://www.ncbi.nlm.nih.gov/sites/GDSbrowser?acc=GDS402). However, the biological role of L13Rik in regulating MC injury remain unknown.

The cell-cycle arrest is a hallmark of MC hypertrophy (Shankland, 1999), and cyclin-dependent kinase inhibitor 1A (CDKN1A), a kind of cyclin-dependent kinase inhibitor, is required for glomerular hypertrophy in experimental DN (Al-Douahji et al., 1999). In addition, up-regulated CDKN1B by lncRNA-NEAT1 promotes MC hypertrophy (Liao et al., 2020). In the present study, we demonstrated that L13Rik was significantly increased in DN patients and rats, and L13Rik depletion effectively reduced HG-induced MC survival, ECM accumulation, and cell hypertrophy through sponging miR-2861 and thus de-repressing its target CDKN1B.

## 2. Materials and Methods

### 2.1 Patients and samples

Human blood specimens were obtained with the approval of the Ethics Committee of Shanghai Jiao Tong University Affiliated Tong Ren Hospital (No. 2019-060) according to the Declaration of Helsinki. In total, 13 DN blood samples were obtained from DN patients aged 53±12 (7 men and 6 women), and 8 control blood samples were obtained from healthy volunteers aged 55±10 (4 men and 4 women) in Tong Ren Hospital. DN was diagnosed in accordance with previous criteria (An et al., 2015). Written informed consent was acquired from each donor before sampling. These donors had not received any therapies within 3 months before blood sampling.

### 2.2 Animal model

All procedures were approved by the Animal Ethics Committee of Shanghai Jiao Tong University Affiliated Tong Ren Hospital and performed under the ARRIVE guidelines. Male Sprague-Dawley (SD) rats, aged 8 weeks, were obtained from Model Animal Research Center of Nanjing University (Nanjing, China), and housed in thermostatic room (25 ± 2 □, 12/12 light/dark cycle, 50% ± 10% relative humidity). Rats were allowed to access chow and water ad libitum. Rats were weighed and randomly divided into control group (n=10) and diabetic group (n=10). Experimental DN was induced on diabetic rats by intraperitoneal injection with streptozotocin (50 mg/kg; Aladdin, Shanghai, China) dissolved in sodium citrate buffer (pH=4.5). Access to 20% glucose water was allowed ad libitum for 18 h to prevent transient hypoglycemia. Seventy-two h after streptozotocin administration, the blood glucose of each rat was measured. Rats with blood glucose > 16.7 mmol/L were considered diabetic. During the experimental period, any rat suffering from disease or dying was euthanized by cervical dislocation under deep anesthesia using pentobarbital sodium (40 mg/kg).

### 2.3 Cell Culture

A murine mesangial cell line, SV40-MES-13, was purchased from American Type Culture Collection (ATCC, MA, USA), and maintained in DMEM (Gibco, CA, USA) containing 10% FBS (Sigma-Aldrich, MO, USA) at 37 □ under 5% CO_2_. To explore the function of glucose concentration on L13Rik expression, MCs were challenged with 30 mM mannitol (Ma) or glucose at concentrations of 5 mM (normal glucose, NG), 15mM, and 30 mM (high glucose, HG). After exposure to glucose, cells were harvested for qRT-PCR and western blot assay.

### 2.4 Overexpression and RNA interference (RNAi)

The recombinant lentivirus harboring full-length L13Rik (Lv-L13Rik) and negative control lentivirus were purchased from Sangon (Shanghai, China). L13Rik siRNA (siL13Rik) and miR-2861 mimics were synthesized by Genewiz (Jiangsu, China), and transfected into MCs with Lipofectamine RNAiMAX (Invitrogen, CA, USA) according to the manufacturer’s introduction. Sequence for RNAi and miRNA sequence was showed in Table 1.

**Table-1.**
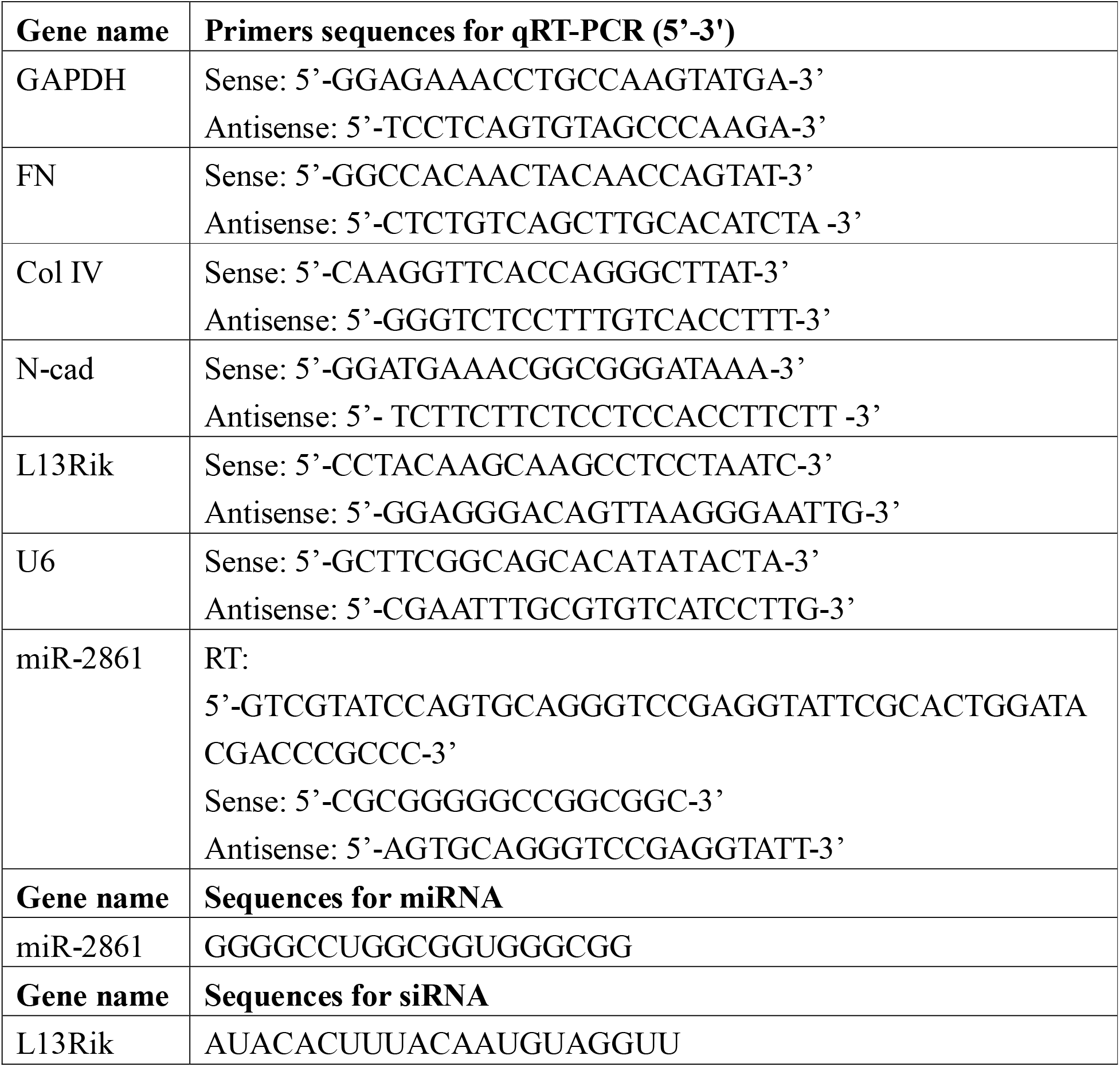
Sequences for qRT-PCR, RNAi, and miRNA.

### 2.5 Cell viability assay

SV40-MES-13 cell viability was assessed using CCK-8 reagent (Beyotime, Shanghai, China) following the manufacturer’s protocols. Briefly, SV40-MES-13 cells were plated into 96-well plates at a density of 3×10^5^ cells per well. SV40-MES-13 cells were cultured in medium containing 30 mM glucose in the presence or absence of miR-2861 mimics or siL13Rik. After culturing for 48 h at 37 □, cells were incubated with 10 μL of CCK-8 reagent for 60 min. The absorbance (450 nm) of each well was measured on a microplate reader (HENGMEI TECHNOLOGY, Shandong, China).

### 2.6 Terminal deoxynucleotidyl transferase (TdT) dUTP nick-end labelling (TUNEL)

SV40-MES-13 cell apoptosis was assessed using TUNEL. In brief, SV40-MES-13 cells were seeded in 12-well plates and cultured in medium containing 30 mM glucose in the presence or absence of miR-2861 mimics or siL13Rik. After incubation for 48 h at 37 □, cells were fixed with 4% PFA (Beyotime) for 20 min and dyed with TUNEL reagent (Beyotime) for 60 min. Nuclei were labeled using DAPI (Beyotime). The fluorescence signal was captured using MF53-M microscope (Mshot, Guangdong, China).

### 2.7 Fluorescence in Situ Hybridization (FISH)

The L13Rik probe was synthesized by Sangon (Shanghai, China). After culturing in 12-well plates in the presence or absence of HG for 48 h, SV40-MES-13 cells were washed by PBS and fixed by 4% PFA for 15 minutes at room temperature, following treatment with proteinase K, glycine and acetic anhydride. Pre-hybridization was performed at 37 □ for 1 h, followed by hybridization L13Rik against using 250 ng/mL of L13Rik probe. Nuclei were labeled with DAPI, and L13Rik signal was captured as mentioned earlier.

### 2.8 qRT-PCR

Total RNA in blood samples, renal tissues, and SV40-MES-13 cells was extracted with RNA simple Total RNA Kit (Tiangen, Beijing, China). cDNA was synthesized with TIANScript□RT Kit (Tiangen) at 42 □ for 60 min. qRT-PCR was performed in triplicate with FastFire qPCR PreMix (SYBR Green) (Tiangen) on ViPlex Fluor Real-time PCR system (Vivantis, Selangor, Malaysia). Thermocycling conditions were set as follows: 95 □ for 60 s, followed by 30 cycles of 95 □ for 15 s and 67□ for 30s. Gene expressions were calculated by 2^-ΔΔCT^ method. Expressions of Fibronectin (FN), collagen IV (Col-IV), N-cadherin (N-cad), CDKN1B and L13Rik were normalized by GAPDH, and expression of miR-2861 was normalized by U6. All primer sequences were listed in Table 1.

### 2.9 Western blot

SV40-MES-13 cells were lysed with xTractor Buffer Kit (CLONTECH Laboratories, Inc., CA, USA). Protein specimens were quantified through BCA method and electrophoresed through 10% SDS-PAGE gels. After transferring onto PVDF membranes, interested proteins were masked with 2% non-fat milk at 37 □ for 30 min, then incubated with anti-Fibronectin (FN) antibody (1:700; SAB5700724; Sigma-Aldrich), anti-collagen IV (Col-IV) antibody (1: 500; ab52235; Abcam, CA, USA), anti-CDKN1B antibody (1: 1000; ab32034; Abcam), anti-N-cadherin (N-cad) antibody (1:8000; ab76011; Abcam), and anti-β-actin antibody (1:20000; AF7018; Affinity, Jiangsu, China) at 4 □ for 20 h. The membranes were rinsed 3 times with PBST, then immersed in HRP-conjugated anti-rabbit IgG secondary antibody solution (1: 5000; S0001; Affinity) at 37 □ for 60 min. Interesting proteins were visualized using commercial ECL kit (Glpbio, CA, USA). Fluorescence intensities were quantified using Fiji software.

### 2.10 RNA pull-down assay

Biotin-labeled miR-2861 was purchased from Genewiz (Jiangsu, China). Biotin-labelled miR-2861 (3 μg) was incubated with streptavidin-coated beads (434341; Thermofisher Scientific) at 4□°C overnight. Then, streptavidin-coated beads were incubated with SV40-MES-13 cells lysate at 4□°C for 12 h. RNAs bound by miR-2861 were purified with Trizol reagent (Takara, Shiga, Japan), and assessed through qRT-PCR.

### 2.11 RNA immunoprecipitation (RIP) assay

After miR-2861 knockdown, RIP assay was performed as per the instructions of the Immunoprecipitation Kit with Protein A+G Magnetic Beads (Beyotime). At first, SV40-MES-13 cells (1×10^6^) were lysed with 200 μL RIPA buffer (Beyotime). Lysates were incubated with 20 μl of magnetic beads labeled with normal mouse IgG (A7028; Beyotime) or rabbit anti-Ago2 antibody (DF12246; Affinity) at 4 □ for 16 h. Immunoprecipitated RNA was obtained by digesting protein with Proteinase-K, and quantified on a microplate reader. Lastly, immunoprecipitated L13Rik and miR-2861 were assessed through qRT-PCR.

### 2.12 Dual□luciferase reporter assay

The wild-type or mutant predictive binding site of miR-2861 on L13Rik was cloned into PGL3 vector (Fenghui Biotechnology, Hunan, China) to construct pGL3-L13Rik-WT or pGL3-L13Rik-Mut plasmids. 3’UTR of CDKN1B or its mutant was inserted into PGL3 vector to construct pGL3-CDKN1B-3’UTR-WT or pGL3-CDKN1B-3’UTR-Mut plasmids. HEK293 cells (1×10^6^) were plated into 24-well plates. 20ng of pGL3-L13Rik-WT, pGL3-L13Rik-Mut, pGL3-CDKN1B-3’UTR-WT or pGL3-CDKN1B-3’UTR-Mut plasmids was co-transfected with 40 nM of miR-2861, and 2 ng of pRL-TK (Beyotime) into HEK293 cells with Lipofectamine™ 3000 according to the manufacturer’s protocol. After incubation at 37 □ for 48 h, luciferase activities were assessed using Dual-Luciferase Reporter Gene Assay Kit (Beyotime) in accordance with the manufacture’s protocol. Renilla luciferase was used for an internal control.

### 2.13 Cell hypertrophy

SV40-MES-13 cell hypertrophy was assessed by measuring the ratio of total protein to cell number, as described previously (Dey et al., 2015).

### 2.14 Statistical analysis

Data were presented as mean ± standard error of the mean (SEM) from three independent experiments and analyzed using SPSS 22.0 software (IBM, USA). The difference among groups was compared with student’s *t*-test or one-way analysis of variance followed by the Scheffé test. P-value less than 0.05 was defined as statistically significant.

## 3. Results

### 3.1 L13Rik was increased in DN patients, diabetic rats, and HG-treated MCs

Given that L13Rik expression is up-regulated in renal tissues of mice with type 2 diabetes compared with normal mice (Supporting Figure S1), we next investigated whether L13Rik is correlated with DN progression through regulating MC injury. To this end, L13Rik level was first assessed in DN patients and diabetic rats. As shown in Figure 1A, L13Rik level was significantly up-regulated in peripheral blood of DN patients compared with the healthy controls. Similarly, L13Rik expression was also increased in renal tissues of diabetic rats compared with control rats (Figure 1B). Consistently, Figure 1C showed that L13Rik expression was remarkably up-regulated in HG-cultured SV40-MES-13 cells. The relationships between treatment time and glucose concentration on L13Rik expression were also explored. As shown in Figure 1D and E, HG resulted in a significant increase of L13Rik level in a time-dependent and dose-dependent manner. Furthermore, the results from FISH assay verified that L13Rik was up-regulated in HG-cultured SV40-MES-13 cells, and that L13Rik was mainly located in the cytoplasm of SV40-MES-13 cells (Figure 1F).

**Figure 1.**
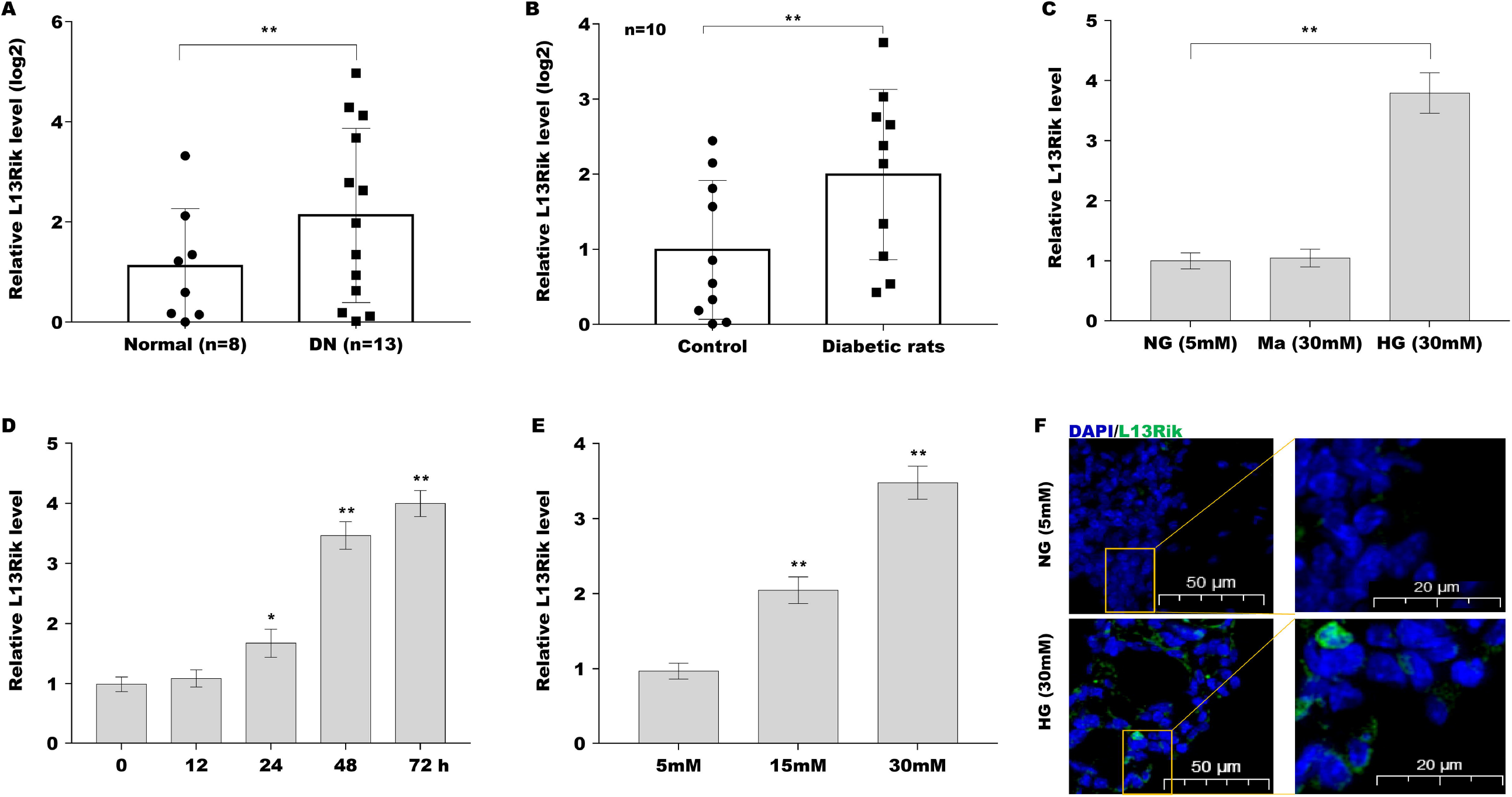
L13Rik was up-regulated in DN patients, diabetic rats, and HG-treated SV40-MES-13 cells. qRT-PCR analysis of L13Rik level in peripheral blood of DN patients (n=13) and healthy controls (n=8) (A), in renal tissues of diabetic rats (n=10) (B), and in HG-cultured SV40-MES-13 cells (C). qRT-PCR analysis of L13Rik level in SV40-MES-13 cells after treatment with 30 mM glucose for different time (D) or treatment with different dose of glucose (E). (F) FISH analysis for L13Rik cellular localization in SV40-MES-13 cells. Probes targeting L13Rik were stained in green and the nucleuses were stained in blue. \**p*<0.05. \*\**p*<0.01.

### 3.2 L13Rik depletion suppressed HG-induced MC survival, ECM accumulation, and hypertrophy

To investigate the biological effect of L13Rik on DN, SV40-MES-13 cells were treated with siRNA against L13Rik (siL13Rik) and then cell injury was assessed through analyzing cell viability and ECM accumulation. The results from CCK-8 assay showed that SV40-MES-13 cell viability was markedly increased after treatment with HG, whereas L13Rik depletion significantly reversed the effect (Figure 2A). Tunel assay showed that SV40-MES-13 cell apoptosis was markedly decreased after treatment with HG, whereas L13Rik depletion significantly reversed the effect (Figure 2B and C).

**Figure 2.**
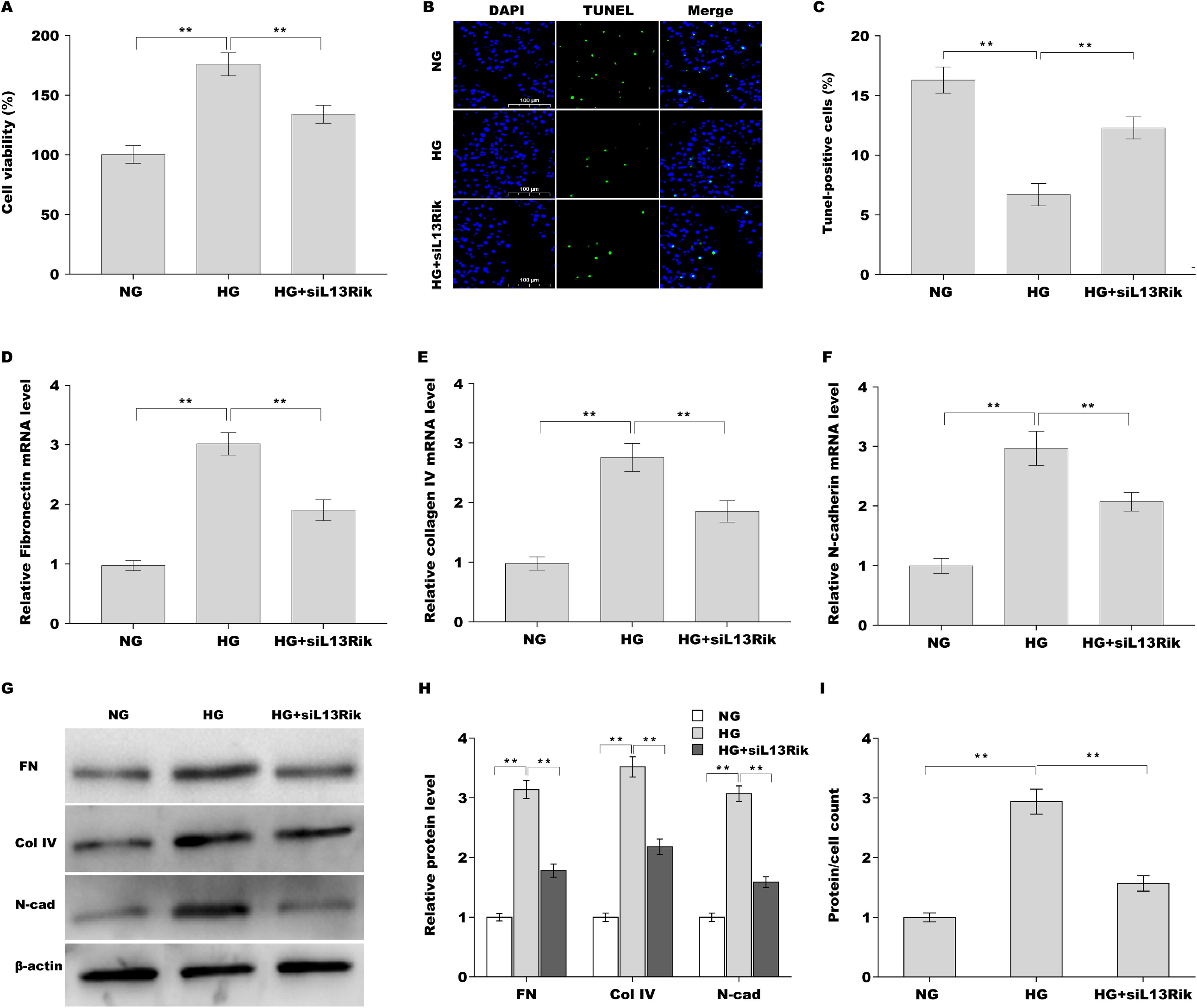
L13Rik depletion suppressed HG-induced MC survival, ECM accumulation, and hypertrophy. (A) SV40-MES-13 cell viability was assessed using CCK-8 after HG treatment with or without L13Rik knockdown. (B) SV40-MES-13 cell apoptosis was assessed using Tunel staining after HG treatment with or without L13Rik knockdown. (C) The mRNA levels of FN, Col-IV, and N-cad in SV40-MES-13 cells were assessed by qRT-PCR after HG treatment with or without L13Rik knockdown. (D and E) Western blot and quantitative analysis of FN, Col-IV, and N-cad protein expression in SV40-MES-13 cells after HG treatment with or without L13Rik knockdown. (F) SV40-MES-13 cell hypertrophy was assessed according to total protein content per cell. \**p*<0.05. \*\**p*<0.01.

To identify the effect of L13Rik on ECM accumulation in SV40-MES-13 cells, FN, Col-IV and N-cad expression was assessed by qRT-PCR and western blot analysis after treatment with HG in the presence or absence of siL13Rik. Figure 2D-F showed that the mRNA levels of FN, Col-IV and N-cad were markedly increased after treatment with HG, whereas the effect was blocked by L13Rik depletion. As expected, the protein levels of FN, Col-IV and N-cad were remarkably increased after treatment with HG, whereas the effect was blocked by L13Rik depletion (Figure 2G and H). Furthermore, L13Rik silencing alleviated HG-induced SV40-MES-13 cell hypertrophy, as suggested by a decreased total protein content per cell (Figure 2I).

### 3.3 L13Rik acted as ceRNA to sponge miR-2861

LncRNA, located in cytoplasm, commonly acts as ceRNA to control gene expression via sponging specific miRNAs (Thomson and Dinger, 2016). Given that L13Rik mainly locates in the cytoplasm in SV40-MES-13 cells (Figure 1F), we speculated L13Rik might exert its biological effect in this way. To prove that, a bioinformatics tool, miRDB (http://mirdb.org/cgi-bin/search.cgi.) (Chen and Wang, 2020), was applied to predict miRNAs sponged by L13Rik, and 63 candidate miRNAs were identified (Supporting Table S1). A previous study has demonstrated that 86 miRNAs were dys-expressed in DN progression (Supporting Table S2) (Chen et al., 2020). Venn diagram analysis revealed that miR-2861 was the only miRNA that could be sponged by L13Rik and dy-sregulated in DN patients (Supporting Figure S3).

MiR-2861 level was next assessed in DN patients and diabetic rats. As shown in Figure 3A, miR-2861 expression was obviously decreased in peripheral blood of DN patients compared with healthy controls. Figure 3B showed that miR-2861 was also down-regulated in renal tissues of diabetic rats compared with control rats. In addition, HG treatment resulted in a significant decrease of miR-2861 in a dose-dependent manner (Figure 3C).

**Figure 3.**
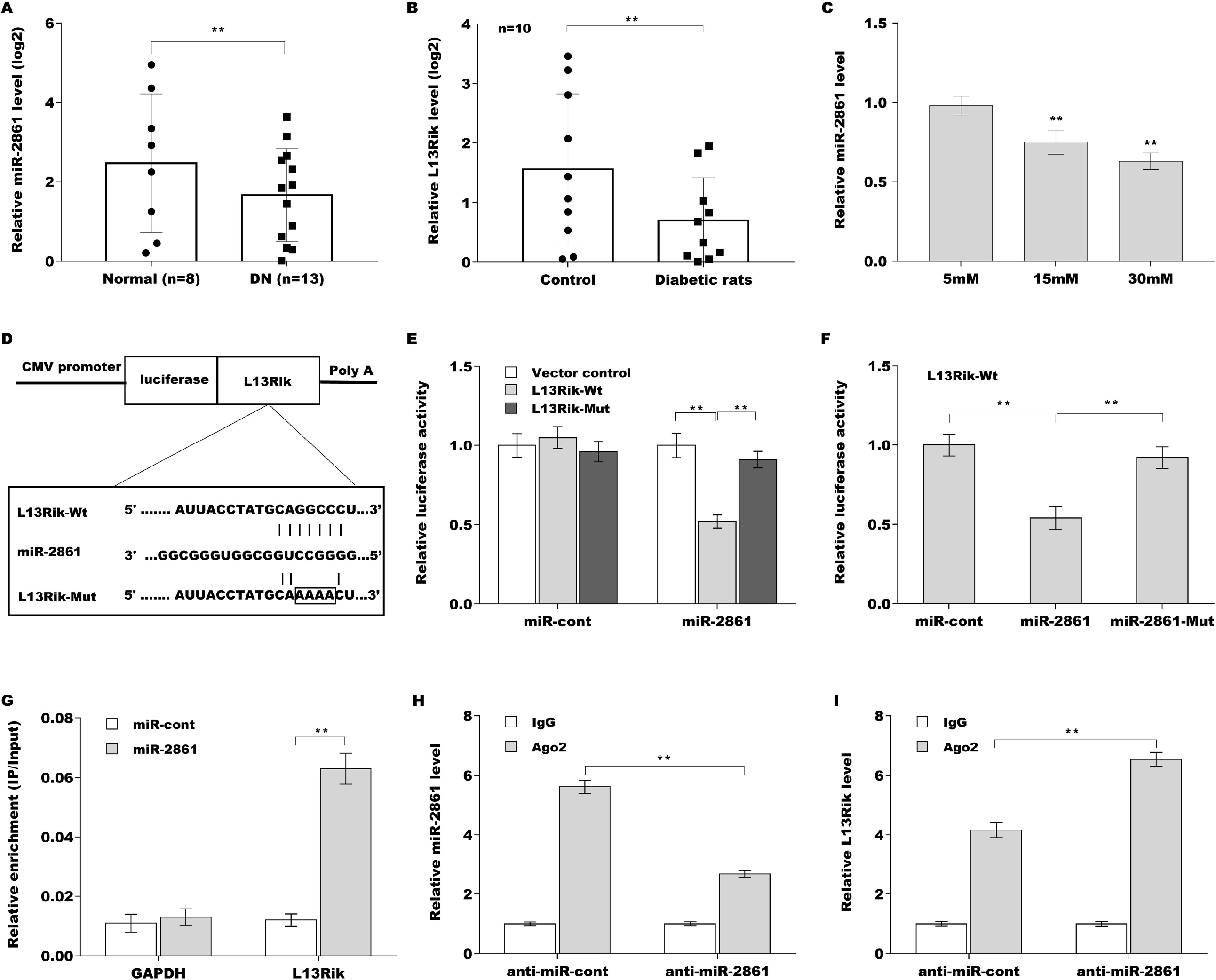
L13Rik sponged miR-2861 by acting as ceRNA. qRT-PCR analysis of miR-2861 level in DN patients (n=13) and 8 healthy controls (n=8) (A), in diabetic rats (n=10) (B), and in HG-cultured SV40-MES-13 cells (C). (D) Schematic diagram of the miR-2861 site in L13Rik-wt-3’UTR and L13Rik-mutant-3’UTR. (E) Luciferase activity was assessed in HEK293 cells co-transfected with miR-2861 and luciferase reporters containing L13Rik-wt-3’UTR and L13Rik-mutant-3’UTR. Data is presented as the relative ratio of firefly luciferase activity to renilla luciferase activity. (F) Luciferase activity was assayed in HEK293 cells co-transfected with L13Rik-wt-3’UTR with miR-2861-wt or miR-2861-Mut. (G) The direct interaction of miR-2861 with L13Rik was assessed by RNA pull-down assay. (H and I) RIP analyses with anti-Ago2 antibody were carried out to assess the enrichment of L13Rik and miR-2861. \**p*<0.05. \*\**p*<0.01.

To verify the direct combination of miR-2861 with L13Rik, the binding sites between L13Rik and miR-2861 were analyzed and recombinant plasmids containing the binding sites (pGL3-L13Rik-Wt and pGL3-L13Rik-Mut) was constructed (Figure 3D). pGL3-L13Rik-Wt or pGL3-L13Rik-Mut was co-transfected with miR-2861 into HEK293 cells and then relative luciferase activity was measured using dual□luciferase reporter assay. As shown in Figure 3E, miR-2861 significantly decreased the luciferase activity of pGL3-L13Rik-Wt, but did not affect the luciferase activity of pGL3-L13Rik-Mut. Furthermore, miR-2861 mutant lost the inhibitor effect to luciferase activity of pGL3-L13Rik-Wt (Figure 3F), indicating that the combination of miR-2861 with L13Rik was sequence specific. The results from RNA pull-down assay further demonstrated that L13Rik was more enriched in the biotin-labelled miR-2861 precipitates than biotin-labelled miR-cont precipitates (Figure 3G). Argonaute 2 (Ago2) is the core component of “miRNA-induced silencing complex (miRISC)”, a multi-protein complex that incorporates miRNA and its target mRNA (or lncRNA) (Chendrimada et al., 2005, Liu et al., 2004). The results from RIP with Ago2 antibody showed that miRNA-2861 and L13Rik were concurrently enriched in SV40-MES-13 cells, and miRNA-2861 inhibitor significantly reduced miRNA-2861 level and increased L13Rik enrichment in Ago2 precipitates (Figure 3H and I). These results demonstrate that L13Rik functions as ceRNA to sponge miR-2861 in a sequence-specific manner.

### 3.4 MiR-2861 suppressed HG-induced MC survival, ECM accumulation, and hypertrophy

The role of miR-2861 in regulating MC injury was next investigated. The results from CCK-8 assay showed that miR-2861 overexpression significantly repressed HG-induced increase of SV40-MES-13 cell viability (Figure 4A). Tunel assay showed that miR-2861 accelerated HG-treated SV40-MES-13 cell apoptosis (Figure 4B). To identify the effect of miR-2861 on ECM accumulation in SV40-MES-13 cells, FN, Col-IV and N-cad expression was assessed by qRT-PCR and western blot analysis after treatment with HG in the presence or absence of miR-2861. Figure 4C showed that miR-2861 significantly repressed HG-induced increase in FN, Col-IV and N-cad mRNA level. miR-2861 also significantly suppressed HG-induced increase in FN, Col-IV and N-cad protein level (Figure 4D and E). Furthermore, miR-2861 alleviated HG-induced SV40-MES-13 cell hypertrophy (Figure 4F).

**Figure 4.**
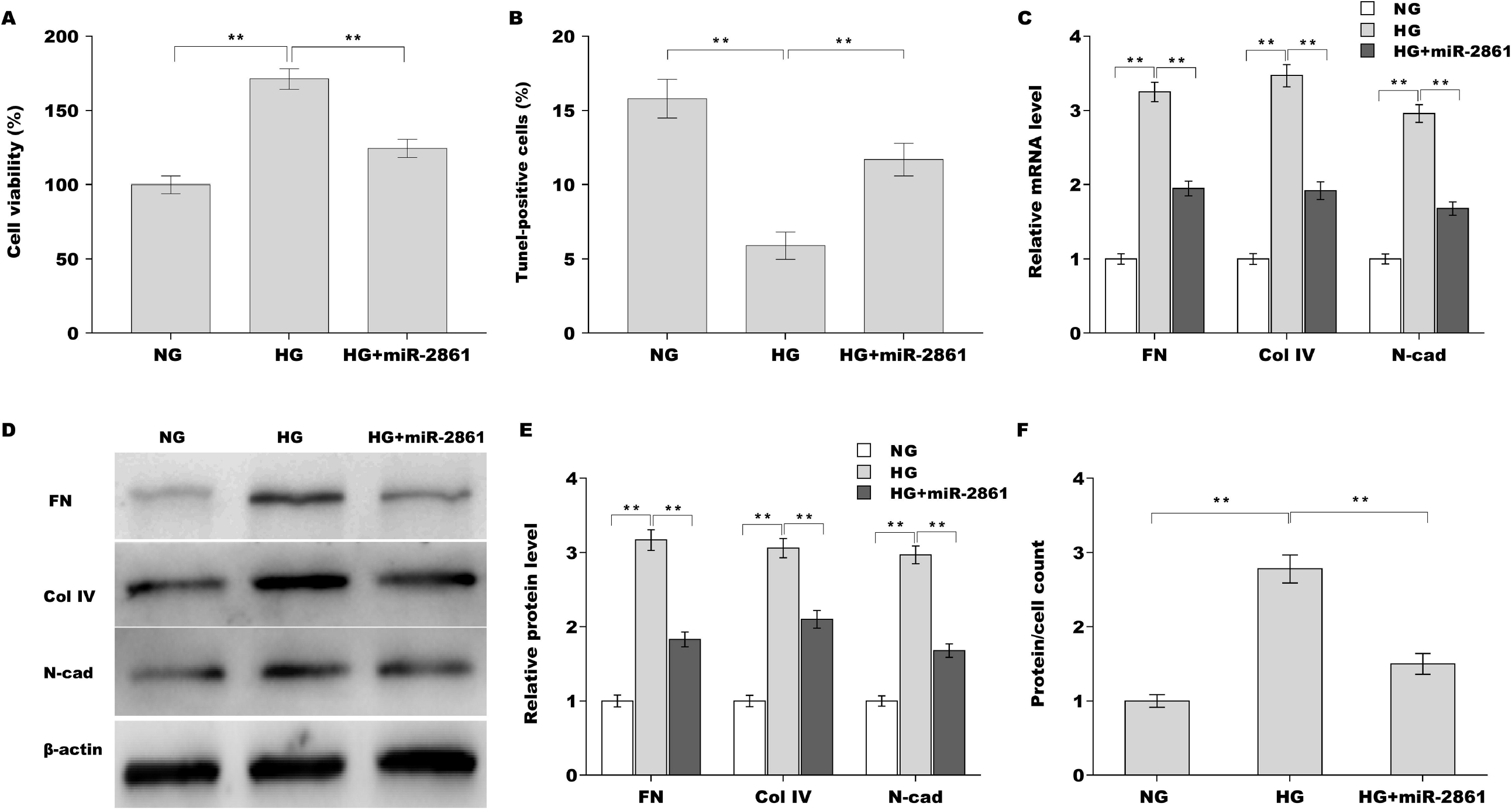
MiR-2861 suppressed HG-induced MC survival, ECM accumulation, and hypertrophy. (A) SV40-MES-13 cell viability was assessed using CCK-8 after HG treatment with or without miR-2861 overexpression. (B) SV40-MES-13 cell apoptosis was assessed using Tunel staining after HG treatment with or without miR-2861 overexpression. (C) The mRNA levels of FN, Col-IV, and N-cad in SV40-MES-13 cells were assessed using qRT-PCR after HG treatment with or without miR-2861 overexpression. (D and E) Western blot and quantitative analysis of FN, Col-IV, and N-cad protein expression in SV40-MES-13 cells after HG treatment with or without miR-2861 overexpression. (F) Cell hypertrophy was assessed using total protein level per cell after HG treatment with or without miR-2861 overexpression. \**p*<0.05. \*\**p*<0.01.

### 3.5 MiR-2861 antagonized the effect of L13Rik on MC injury

To explore whether L13Rik accelerated MC injury through sponging miR-2861, SV40-MES-13 cells were over-expressed by L13Rik in the presence or absence of miR-2861 mimic, and cell viability and EMC production were assessed. As shown in Figure 5A, L13Rik increased SV40-MES-13 cell viability, whereas miR-2861 over-expression reversed the effect. Moreover, L13Rik increased the mRNA and protein expression of FN, Col-IV, and N-cad, whereas miR-2861 over-expression reversed the effect (Figure 5B-D). miR-2861 also repressed L13Rik-induced SV40-MES-13 cell hypertrophy (Figure 5E).

**Figure 5.**
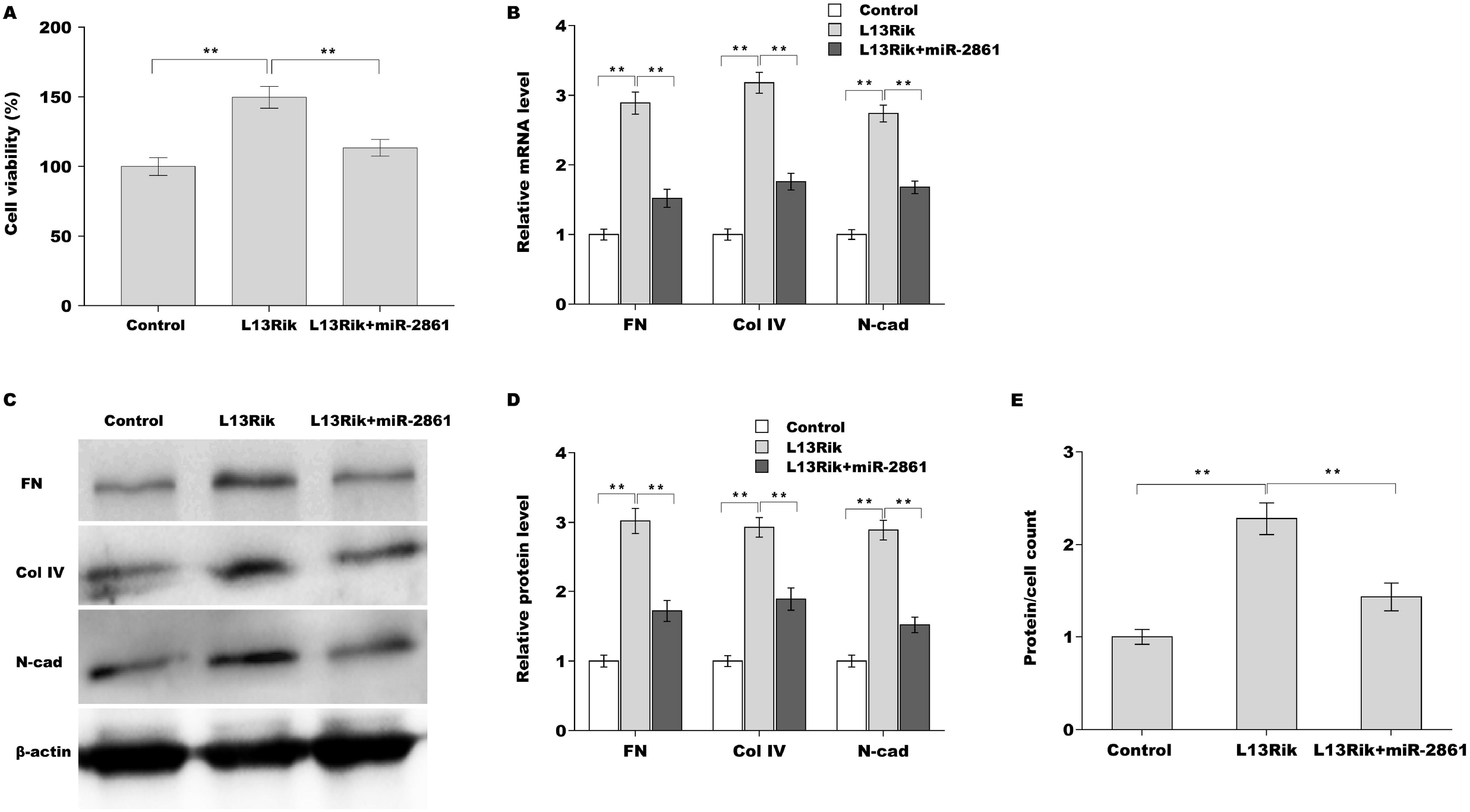
MiR-2861 antagonized the effect of L13Rik on MC injury. (A) SV40-MES-13 cell viability was assessed using CCK-8 after overexpression with L13Rik in the presence or absence of miR-2861. (B) The mRNA levels of FN, Col-IV, and N-cad in SV40-MES-13 cells were assessed by qRT-PCR after overexpression with L13Rik in the presence or absence of miR-2861. (C and D) Western blot and quantitative analysis of FN, Col-IV, and N-cad protein expression in SV40-MES-13 cells after overexpression with L13Rik in the presence or absence of miR-2861. (E) Cell hypertrophy was assessed using total protein level per cell after overexpression with L13Rik in the presence or absence of miR-2861. \*\**p*<0.01.

### 3.6 L13Rik increased CDKN1B expression by sponging miR-2861

The potential target genes of miR-2861 were predicted using TargetScan tool (http://www.targetscan.org/vert_71/). Total 4407 genes were predicted possibly targeted by miR-2861. Among these genes, CDKN1B was previously reported to be associated with MC hypertrophy (Awazu et al., 2003). To reveal the regulatory role of miR-2861 in CDKN1B, recombinant plasmids of pGL3-CDKN1B-3’UTR-Wt and pGL3-CDKN1B-3’UTR-Mut were constructed and co-transfected with miR-2861 (Figure 6A). As shown in Figure 6B, miR-2861 significantly decreased the luciferase activity of pGL3-CDKN1B-3’UTR-Wt but had no impact on luciferase activity of pGL3-CDKN1B-3’UTR-Mut. Moreover, CDKN1B protein expression was further assessed in SV40-MES-13 cells after treatment with miR-2861 mimic. Figure 6C and D showed that CDKN1B expression was obviously decreased after miR-2861 overexpression. More important, L13Rik increased CDKN1B expression in SV40-MES-13 cells, whereas miR-2861 over-expression reversed the effect (Figure 6E and F).

**Figure 6.**
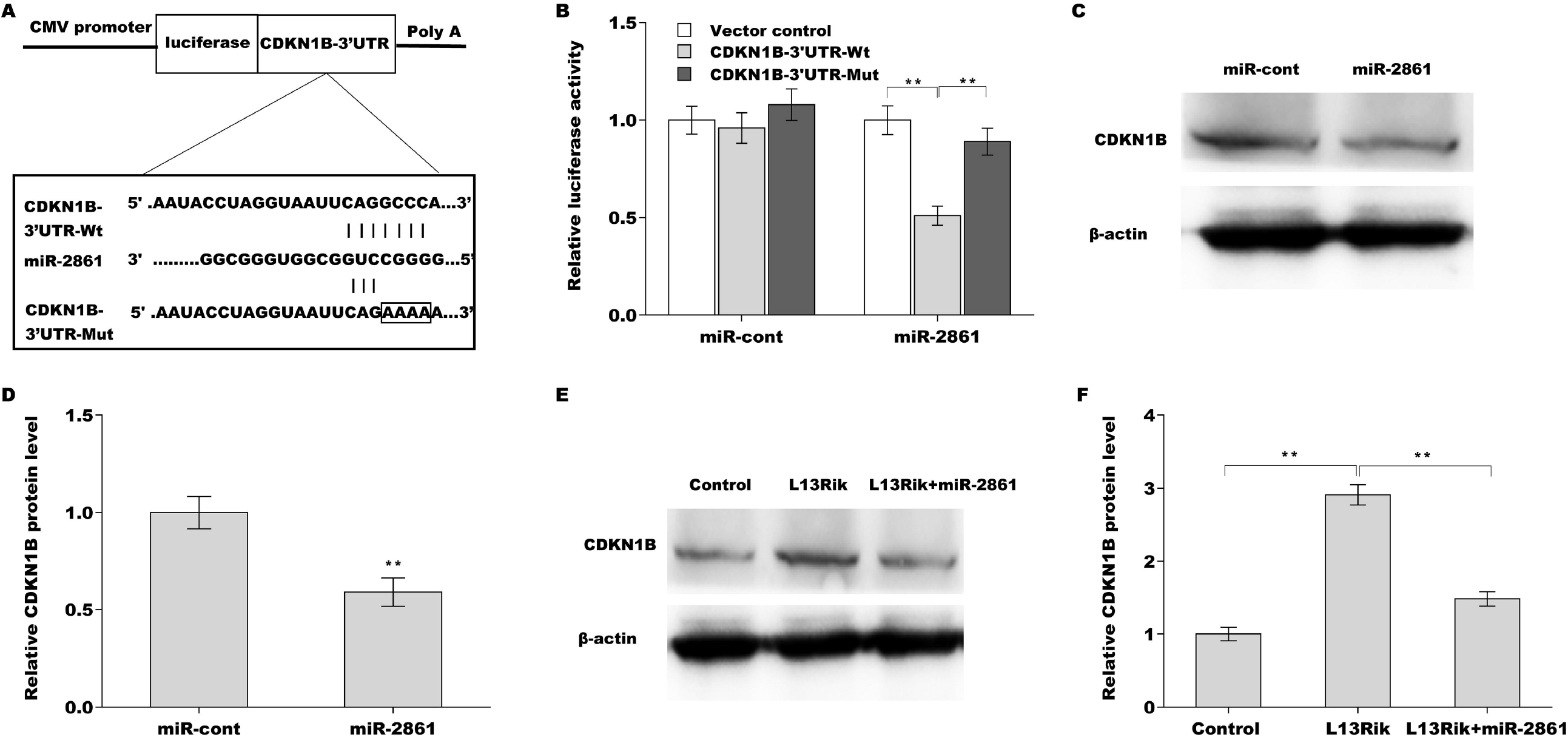
L13Rik promoted MC injury by modulating miR-2861/CDKN1B axis. (A) Schematic diagram of the miR-2861 site in CDKN1B-3’UTR. (B) Luciferase activity was assessed in HEK293 cells co-transfected with miR-2861 and luciferase reporters containing CDKN1B-3’UTR-Wt or CDKN1B-3’UTR-Mut. (C and D) Western blot and quantitative analysis of CDKN1B protein expression in SV40-MES-13 cells after miR-2861 overexpression. (E and F) Western blot and quantitative analysis of CDKN1B protein expression in SV40-MES-13 cells after L13Rik overexpression in the presence or absence of miR-2861. \**p*<0.05. \*\**p*<0.01.

## 4. Discussion

MC injury, including hyperplasia, ECM production, and hypertrophy, plays a crucial role in the pathology of DN (Dai et al., 2017). LncRNA was reported to be involved in MC injury and DN progression (Coellar et al., 2021). However, the role of lncRNA in these processes is still unclear. In the current study, we demonstrated that L13Rik mediated HG-induced MC injury by regulating miR-2861/CDKN1B axis, as evidenced by the following: i) L13Rik was up-regulated in DN patients, diabetic rats and HG-treated MCs; ii) Knockdown of L13Rik suppressed HG-induced MC viability, ECM accumulation and hypertrophy; iii) L13Rik acted as ceRNA to sponge miR-2861; iv) MiR-2861 suppressed HG-induced MC viability, ECM accumulation, and hypertrophy; v) MiR-2861 antagonized the effect of L13Rik on MC injury; vi) L13Rik accelerated MC injury by modulating miR-2861/CDKN1B axis. These data revealed the function and underlying mechanism of L13Rik on regulating MC injury, and may provide a potential opportunity to treat DN.

L13Rik is an lncRNA of 1726 bases in length, which locates in chromosome 3 (Chr3: 95871521-95889093). The abnormal expression of L13Rik has been revealed in HG or TGF-β-treated MCs (GSE2557 and GSE2558) and renal tissues from type 2 diabetic mice (GSE642) using RNA sequencing and microarray analysis. However, the biological role of L13Rik in DN progression remains unknown up to now. In the study we demonstrated that up-regulated L13Rik exerts a significant role in the progress of HG-triggered MC viability, EMC accumulation, and cell hypertrophy. L13Rik depletion remarkably alleviates HG-induced MC injury. Given that kidney is mainly composed of three types of cells: MC, podocyte, and endothelial cell (Wallace, 1998), and the significant role of L13Rik in MC injury, it is essential to explored whether L13Rik regulates HG-induced podocyte, and endothelial cell injury. In an addition, the effect of L13Rik on DN progression will be investigated in diabetic rats. Mounting studies have demonstrated that lncRNA functions as important mediator in DN or other diabetic complications (Lu et al., 2021). For instance, lncRNA MSC-AS1 is up-regulated in peripheral blood from DN patients, and MSC-AS1 silencing alleviates HG-triggered MC proliferation, fibrosis, and inflammation through sponging miR-325 and de-repressing cyclin G1 (Zhao et al., 2022). LncRNA NEAT1 expression is increased in DN patients and HG-treated MCs, and NEAT1 over-expression accelerates MC hypertrophy through sponging miR-222-3p and thus increasing CDKN1B expression (Liao et al., 2020). As a kind of decoys that compete for miRNAs, ceRNA incorporates non-coding RNA (miRNA and lncRNA) with mRNAs in miRISC through miRNA response elements (MREs) (Karreth and Pandolfi, 2013). According to this hypothesis, lncRNAs containing MREs can segregate miRNAs from mRNAs containing the same MREs, and de-repressing the mRNA expression (Karreth and Pandolfi, 2013, Guo et al., 2010). In fact, more and more studies have revealed that lncRNAs function as its biological role in this way. In the study we demonstrated that L13Rik acts as a ceRNA to sponge miR-2861, resulting in the de-repression of its target CDKN1B, a gene known to accelerate MC injury (Liao et al., 2020).

A previous study found that miR-2861 is decreased in DN patients and associated with estimated glomerular filtration rate (Cardenas-Gonzalez et al., 2017). However, the mechanism of miR-2861 in the development of DN is still unclear. The current results revealed that miR-2861 is decreased in DN patients and HG-induced MCs. Functionally, miR-2861 over-expression represses HG-induced MC viability, ECM accumulation, and hypertrophy. In summary, the L13Rik/miR-2861/CDKN1B axis plays an important regulatory role in DN progression.

## Ethics approval and consent to participate

All procedures performed in this study were in accordance with the ethical standards of the institutional and international research committee and with the 1964 Helsinki declaration and its later amendments or comparable ethical standards. ShangHai Tongren Hospital (2019-060, AF/SC-11/D2.1)

## Consent for publication

Informed consent was obtained from all participants included in the study.

## Data Availability

The information generated and analyzed during the current study is available from the corresponding author on reasonable request.

## Supplementary material

Supplementary material is available on the publisher’s website along with the published article.

## Reference

Al-Douahji, M., Brugarolas, J., Brown, P. A., Stehman-Breen, C. O., Alpers, C. E. & Shankland, S. J. 1999. The cyclin kinase inhibitor p21WAF1/CIP1 is required for glomerular hypertrophy in experimental diabetic nephropathy. Kidney Int, 56, 1691–9.

Alicic, R. Z., Rooney, M. T. & Tuttle, K. R. 2017. Diabetic Kidney Disease: Challenges, Progress, and Possibilities. Clin J Am Soc Nephrol, 12, 2032–2045.

An, Y., Xu, F., Le, W., Ge, Y., Zhou, M., Chen, H., Zeng, C., Zhang, H. & Liu, Z. 2015. Renal histologic changes and the outcome in patients with diabetic nephropathy. Nephrol Dial Transplant, 30, 257–66.

Awazu, M., Omori, S., Ishikura, K., Hida, M. & Fujita, H. 2003. The lack of cyclin kinase inhibitor p27(Kip1) ameliorates progression of diabetic nephropathy. J Am Soc Nephrol, 14, 699–708.

Bach, D. H. & Lee, S. K. 2018. Long noncoding RNAs in cancer cells. Cancer Lett, 419, 152–166.

Cardenas-Gonzalez, M., Srivastava, A., Pavkovic, M., Bijol, V., Rennke, H. G., Stillman, I. E., Zhang, X., Parikh, S., Rovin, B. H., Afkarian, M., De Boer, I. H., Himmelfarb, J., Waikar, S. S. & Vaidya, V. S. 2017. Identification, Confirmation, and Replication of Novel Urinary MicroRNA Biomarkers in Lupus Nephritis and Diabetic Nephropathy. Clin Chem, 63, 1515–1526.

Chang, J., Yu, Y., Fang, Z., He, H., Wang, D., Teng, J. & Yang, L. 2020. Long non-coding RNA CDKN2B-AS1 regulates high glucose-induced human mesangial cell injury via regulating the miR-15b-5p/WNT2B axis. Diabetol Metab Syndr, 12, 109.

Chen, L., Wu, B., Wang, S., Xiong, Y., Zhou, B., Cheng, X., Zhou, T., Luo, R., Lam, T. W., Yan, B. & Chen, J. 2020. Identification of Cooperative Gene Regulation Among Transcription Factors, LncRNAs, and MicroRNAs in Diabetic Nephropathy Progression. Front Genet, 11, 1008.

Chen, Y. & Wang, X. 2020. miRDB: an online database for prediction of functional microRNA targets. Nucleic Acids Res, 48, D127–D131.

Chendrimada, T. P., Gregory, R. I., Kumaraswamy, E., Norman, J., Cooch, N., Nishikura, K. & Shiekhattar, R. 2005. TRBP recruits the Dicer complex to Ago2 for microRNA processing and gene silencing. Nature, 436, 740–4.

Choudhury, D., Tuncel, M. & Levi, M. 2010. Diabetic nephropathy -- a multifaceted target of new therapies. Discov Med, 10, 406–15.

Coellar, J. D., Long, J. & Danesh, F. R. 2021. Long Noncoding RNAs and Their Therapeutic Promise in Diabetic Nephropathy. Nephron, 145, 404–414.

Dai, H., Liu, Q. & Liu, B. 2017. Research Progress on Mechanism of Podocyte Depletion in Diabetic Nephropathy. J Diabetes Res, 2017, 2615286.

De Oliveira, J. C., Oliveira, L. C., Mathias, C., Pedroso, G. A., Lemos, D. S., Salviano-Silva, A., Jucoski, T. S., Lobo-Alves, S. C., Zambalde, E. P., Cipolla, G. A. & Gradia, D. F. 2019. Long non-coding RNAs in cancer: Another layer of complexity. J Gene Med, 21, e3065.

Dey, N., Bera, A., Das, F., Ghosh-Choudhury, N., Kasinath, B. S. & Choudhury, G. G. 2015. High glucose enhances microRNA-26a to activate mTORC1 for mesangial cell hypertrophy and matrix protein expression. Cell Signal, 27, 1276–85.

Guo, H., Ingolia, N. T., Weissman, J. S. & Bartel, D. P. 2010. Mammalian microRNAs predominantly act to decrease target mRNA levels. Nature, 466, 835–40.

Jiang, Z. F. & Zhang, L. 2021. LncRNA: A Potential Research Direction in Intestinal Barrier Function. Dig Dis Sci, 66, 1400–1408.

Karreth, F. A. & Pandolfi, P. P. 2013. ceRNA cross-talk in cancer: when ce-bling rivalries go awry. Cancer Discov, 3, 1113–21.

Li, A., Peng, R., Sun, Y., Liu, H., Peng, H. & Zhang, Z. 2018. LincRNA 1700020I14Rik alleviates cell proliferation and fibrosis in diabetic nephropathy via miR-34a-5p/Sirt1/HIF-1alpha signaling. Cell Death Dis, 9, 461.

Li, Y., Zheng, L. L., Huang, D. G., Cao, H., Gao, Y. H. & Fan, Z. C. 2020. LNCRNA CDKN2B-AS1 regulates mesangial cell proliferation and extracellular matrix accumulation via miR-424-5p/HMGA2 axis. Biomed Pharmacother, 121, 109622.

Liao, L., Chen, J., Zhang, C., Guo, Y., Liu, W., Duan, L., Liu, Z., Hu, J. & Lu, J. 2020. LncRNA NEAT1 Promotes High Glucose-Induced Mesangial Cell Hypertrophy by Targeting miR-222-3p/CDKN1B Axis. Front Mol Biosci, 7, 627827.

Liu, J., Carmell, M. A., Rivas, F. V., Marsden, C. G., Thomson, J. M., Song, J. J., Hammond, S. M., Joshua-Tor, L. & Hannon, G. J. 2004. Argonaute2 is the catalytic engine of mammalian RNAi. Science, 305, 1437–41.

Lu, X., Tan, Q., Ma, J., Zhang, J. & Yu, P. 2021. Emerging Role of LncRNA Regulation for NLRP3 Inflammasome in Diabetes Complications. Front Cell Dev Biol, 9, 792401.

Okazaki, Y., Furuno, M., Kasukawa, T., Adachi, J., Bono, H., Kondo, S., Nikaido, I., Osato, N., Saito, R., Suzuki, H., Yamanaka, I., Kiyosawa, H., Yagi, K., Tomaru, Y., Hasegawa, Y., Nogami, A., Schonbach, C., Gojobori, T., Baldarelli, R., Hill, D. P., Bult, C., Hume, D. A., Quackenbush, J., Schriml, L. M., Kanapin, A., Matsuda, H., Batalov, S., Beisel, K. W., Blake, J. A., Bradt, D., Brusic, V., Chothia, C., Corbani, L. E., Cousins, S., Dalla, E., Dragani, T. A., Fletcher, C. F., Forrest, A., Frazer, K. S., Gaasterland, T., Gariboldi, M., Gissi, C., Godzik, A., Gough, J., Grimmond, S., Gustincich, S., Hirokawa, N., Jackson, I. J., Jarvis, E. D., Kanai, A., Kawaji, H., Kawasawa, Y., Kedzierski, R. M., King, B. L., Konagaya, A., Kurochkin, I. V., Lee, Y., Lenhard, B., Lyons, P. A., Maglott, D. R., Maltais, L., Marchionni, L., Mckenzie, L., Miki, H., Nagashima, T., Numata, K., Okido, T., Pavan, W. J., Pertea, G., Pesole, G., Petrovsky, N., Pillai, R., Pontius, J. U., Qi, D., Ramachandran, S., Ravasi, T., Reed, J. C., Reed, D. J., Reid, J., Ring, B. Z., Ringwald, M., Sandelin, A., Schneider, C., Semple, C. A., Setou, M., Shimada, K., Sultana, R., Takenaka, Y., Taylor, M. S., Teasdale, R. D., Tomita, M., Verardo, R., Wagner, L., Wahlestedt, C., Wang, Y., Watanabe, Y., Wells, C., Wilming, L. G., Wynshaw-Boris, A., Yanagisawa, M., Yang, I., Yang, L., Yuan, Z., Zavolan, M., Zhu, Y., Zimmer, A., Carninci, P., Hayatsu, N., Hirozane-Kishikawa, T., Konno, H., Nakamura, M., Sakazume, N., Sato, K., Shiraki, T., Waki, K., Kawai, J., Aizawa, K., Arakawa, T., Fukuda, S., Hara, A., Hashizume, W., Imotani, K., Ishii, Y., Itoh, M., Kagawa, I., Miyazaki, A., Sakai, K., Sasaki, D., Shibata, K., Shinagawa, A., Yasunishi, A., Yoshino, M., Waterston, R., Lander, E. S., Rogers, J., Birney, E. & Hayashizaki, Y. 2002. Analysis of the mouse transcriptome based on functional annotation of 60,770 full-length cDNAs. Nature, 420, 563–73.

Rojas-Canales, D. M., Li, J. Y., Makuei, L. & Gleadle, J. M. 2019. Compensatory renal hypertrophy following nephrectomy: When and how? Nephrology (Carlton), 24, 1225–1232.

Sakatani, T., Kaneda, A., Iacobuzio-Donahue, C. A., Carter, M. G., De Boom Witzel, S., Okano, H., Ko, M. S., Ohlsson, R., Longo, D. L. & Feinberg, A. P. 2005. Loss of imprinting of Igf2 alters intestinal maturation and tumorigenesis in mice. Science, 307, 1976–8.

Selby, N. M. & Taal, M. W. 2020. An updated overview of diabetic nephropathy: Diagnosis, prognosis, treatment goals and latest guidelines. Diabetes Obes Metab, 22 Suppl 1, 3–15.

Shankland, S. J. 1999. Cell cycle regulatory proteins in glomerular disease. Kidney Int, 56, 1208–15.

Sun, H. J. 2019. Current Opinion for Hypertension in Renal Fibrosis. Adv Exp Med Biol, 1165, 37–47.

Thomson, D. W. & Dinger, M. E. 2016. Endogenous microRNA sponges: evidence and controversy. Nat Rev Genet, 17, 272–83.

Wallace, M. A. 1998. Anatomy and physiology of the kidney. AORN J, 68, 800, 803-16, 819-20; quiz 821-4.

Zhang, P., Sun, Y., Peng, R., Chen, W., Fu, X., Zhang, L., Peng, H. & Zhang, Z. 2019. Long non-coding RNA Rpph1 promotes inflammation and proliferation of mesangial cells in diabetic nephropathy via an interaction with Gal-3. Cell Death Dis, 10, 526.

Zhao, H., Cui, Y., Dong, F. & Li, W. 2022. lncRNA MSC-AS1 Aggravates Diabetic Nephropathy by Regulating the miR-325/CCNG1 Axis. J Healthc Eng, 2022, 2279072.

